# Dopamine D1 receptor availability is not associated with delusional ideation measures of psychosis proneness

**DOI:** 10.1101/321646

**Authors:** Granville James Matheson, Pontus Plavén-Sigray, Anaïs Louzolo, Jacqueline Borg, Lars Farde, Predrag Petrovic, Simon Cervenka

**Affiliations:** Department of Clinical Neuroscience, Center for Psychiatry Research, Karolinska Institutet and Stockholm County Council, SE-171 76 Stockholm, Sweden; Department of Clinical Neuroscience, Karolinska Institutet, SE-171 76 Stockholm, Sweden; PET Science Centre, Precision Medicine and Genomics, IMED Biotech Unit, AstraZeneca, Karolinska Institutet, Sweden

## Abstract

The dopamine D1 receptor (D1R) is thought to play a role in psychosis and schizophrenia, however the exact nature of this involvement is not clear. Positron emission tomography studies comparing D1R between patients and control subjects have produced inconsistent results. An important confounding factor in most clinical studies is previous exposure to antipsychotic treatment, which is thought to influence the density of D1R. To circumvent some of the limitations of clinical studies, an alternative approach for studying the relationship between D1R and psychosis is to examine individuals at increased risk for psychotic disorders, or variation in subclinical psychotic symptoms such as delusional ideation within the general population, referred to as psychosis proneness traits. In this study, we investigated whether D1R availability is associated with delusional ideation in healthy controls using data from 76 individuals measured with PET using [^11^C]SCH23390 and 217 individuals who completed delusional ideation questionnaires, belonging to three different study cohorts. We first performed exploratory, hypothesis-generating, analyses by creating and evaluating a new measure of delusional ideation (n=132 and n=27), which was then found to show a negative association with D1R availability (n=24). Next, we performed confirmatory analyses using Bayesian statistical modelling, in which we first attempted to replicate this result (n=20), and then evaluated the association of Peters Delusion Inventory scores with D1R availability in two independent cohorts (n=41 and 20). Collectively, we found strong evidence that there is little to no linear association between delusional ideation and D1R availability in healthy controls. If differences in D1R can be confirmed in drug-naive schizophrenia patients compared to controls, further studies are needed to ascertain whether these changes occur at the onset of psychotic symptoms or if they are associated with specific behavioural or genetic aspects of psychosis proneness other than delusional ideation.

## Introduction

The dopamine system has been centrally implicated in the pathophysiology of schizophrenia for over fifty years, due primarily to the fact that antipsychotic drugs exert their effect by blocking D2 dopamine receptor (D2R) (Farde, Wiesel, Halldin, & Sedvall, 1988; Nordström et al., 1993; Kapur & Mamo, 2003). In vivo molecular imaging studies using positron emission tomography (PET) have provided a wealth of evidence for elevations in presynaptic dopamine synthesis capacity and amphetamine-induced dopamine release in schizophrenia (Howes & Kapur, 2009). Importantly, increases in presynaptic dopamine synthesis capacity have also been observed in individuals at high risk of developing schizophrenia (Howes et al., 2009), with greater levels observed specifically for those individuals who later developed a psychotic disorder compared to those who did not (Howes et al., 2011). These observations suggest that changes in dopamine function are evident prior to the onset of psychosis.

With regard to dopamine receptors, there have been a series of PET studies since the 1980s examining striatal D2 receptor availability in schizophrenia, finding evidence for a small increases in patients compared to controls (Howes et al., 2012). In contrast, there have only been a handful of PET studies examining the D1 receptor (D1R) in schizophrenia. Compared with the D2R, there is a high concentration of D1R in the cortex (Hall et al., 1994). The frontal cortex, and the dorsolateral prefrontal cortex (DLPFC) in particular, is thought to be a crucial brain region for understanding the biological underpinnings of schizophrenia symptoms (Wagstyl et al., 2016; Cannon et al., 2002; Selemon & Goldman-Rakic, 1999; Callicott et al., 2000), and further study of the D1R in this region therefore has the potential to provide important insights into the postsynaptic dopaminergic aberrations which may occur in the disorder.

In-vivo studies of the D1R performed thus far in schizophrenia patients have shown mixed results. Initial studies using [^11^C]NNC112 and [^11^C]SCH23390 found lower (Okubo et al., 1997), higher (Abi-Dargham et al., 2002), and no difference (Karlsson, Farde, Halldin, & Sedvall, 2002) in the availability of D1R in frontal cortex compared to healthy control subjects. In follow-up studies, the former two research groups both replicated their own respective results in chronic, medicated (henceforth implied by chronic) (Kosaka et al., 2010), and a subsample of drug naive (Abi-Dargham et al., 2012) schizophrenia patients respectively. Lastly, in a small sample of twin pairs discordant for schizophrenia, (Hirvonen et al., 2006) reported decreases in D1R binding in probands with chronic schizophrenia compared to controls. In contrast, increases were shown in monozygotic unaffected co-twins, i.e. individuals at high genetic risk for the disease. Importantly, in studies where both drug naive and medicated patients were examined, the latter group has consistently exhibited numerically lower D1R binding (Okubo et al., 1997; Abi-Dargham et al., 2002; Abi-Dargham et al., 2012; Poels, Girgis, Thompson, Slifstein, & Abi-Dargham, 2013). This may be partly explained by a reduction in D1R as a consequence of antipsychotic medication, which was shown in experimental studies in non-human primates (NHPs) (Lidow & Goldman-Rakic, 1994; Lidow, Elsworth, & Goldman-Rakic, 1997) (although see (Knable, Hyde, Murray, Herman, & Kleinman, 1996)). Although all of the patient studies have been conducted with small sample sizes, and therefore low statistical power, a tentative interpretation of the results is that drug-naive patients with psychosis disorders, as well as unmedicated individuals at high genetic risk for schizophrenia, show increases in frontal cortex D1R binding (Cervenka, 2018). Additional research is needed to understand the role of the D1R in schizophrenia, especially in the early stages of the illness or even prior to its onset in psychosis-prone individuals.

Core symptoms of psychosis are hallucinations (false perceptions) and delusions (irrational or strange inflexible beliefs), referred to as positive symptoms. It has been shown that delusional beliefs (Peters, Joseph, Day, & Garety, 2004; Peters, Joseph, & Garety, 1999), hallucinations and other anomalous perceptions (Bell, Halligan, & Ellis, 2005) are not uncommon in normal populations of whom the vast majority never go on to develop a psychotic disorder. Moreover, delusional beliefs and anomalous perceptions appear to co-vary to a large extent (Bell, Halligan, & Ellis, 2005), are more commonly experienced by relatives of schizophrenia patients than by the general population (Schürhoff et al., 2003), and may constitute an important risk factor for later psychosis (Freeman, 2006; van Os, Linscott, Myin-Germeys, Delespaul, & Krabbendam, 2008). They are thus conceptualised as psychosis proneness traits, whose underlying cognitive, thought- and perceptual mechanisms are considered to be similar to those associated with clinical psychotic symptoms (Schmack et al., 2013; Schmack et al., 2015; Teufel et al., 2015; Powers, Mathys, & Corlett, 2017). Studying these traits in healthy populations allows for the examination of the biological correlates of individual psychotic-like symptoms, without the confounding effect of medication or other limitations common in clinical samples such as comorbidity, disease heterogeneity or differences in physical or cognitive health between groups.

Here, we aimed to investigate the association between D1R availability and delusional ideation in three cohorts of healthy control subjects. Given the evidence for increases in D1R in drug-naive schizophrenia patients (Abi-Dargham et al., 2012), and individuals at high genetic risk for schizophrenia (Hirvonen et al., 2006), we hypothesised that there would be a positive association between delusional ideation and D1R binding in the DLPFC. First, we constructed a new scale for delusional ideation based on a commonly used personality scale. After evaluating its reliability and convergent validity, we performed an exploratory study to investigate the relationship between this scale and D1R availability using previously collected data. We then performed three confirmatory studies to examine the relationship between D1R availability and delusional ideation using both the new scale and the Peters Delusion Inventory (PDI) (Peters, Joseph, Day, & Garety, 2004) in two independent cohorts.

## Methods

### Study design

This investigation consists of four substudies from four separate data collections. A summary of the substudies is presented in Figure 1. Substudies 1A-C were exploratory (i.e. hypothesis-generating), while Substudies 2-4 were confirmatory (i.e. hypothesis-testing) studies. Our investigation was divided in this way in order to ensure that we could explore the data fully and transparently, and subsequently commit to a principled test of our hypotheses based on previous results from our own exploratory findings as well as the scientific literature. This avoids the potential for presenting the conclusions of our experimental studies as if they were confirmatory based on hindsight bias, and ensures that the statistics from our confirmatory studies are valid, and hence possess greater evidentiary weight (Wagenmakers, Wetzels, Borsboom, van, & Kievit, 2012). We consider Substudies 2-4 to be confirmatory despite not being pre-registered (Wagenmakers, Wetzels, Borsboom, van, & Kievit, 2012; Nosek, Ebersole, DeHaven, & Mellor, 2018), due to our use of predictions stemming from the scientific literature and from the exploratory studies.

**Figure 1:**
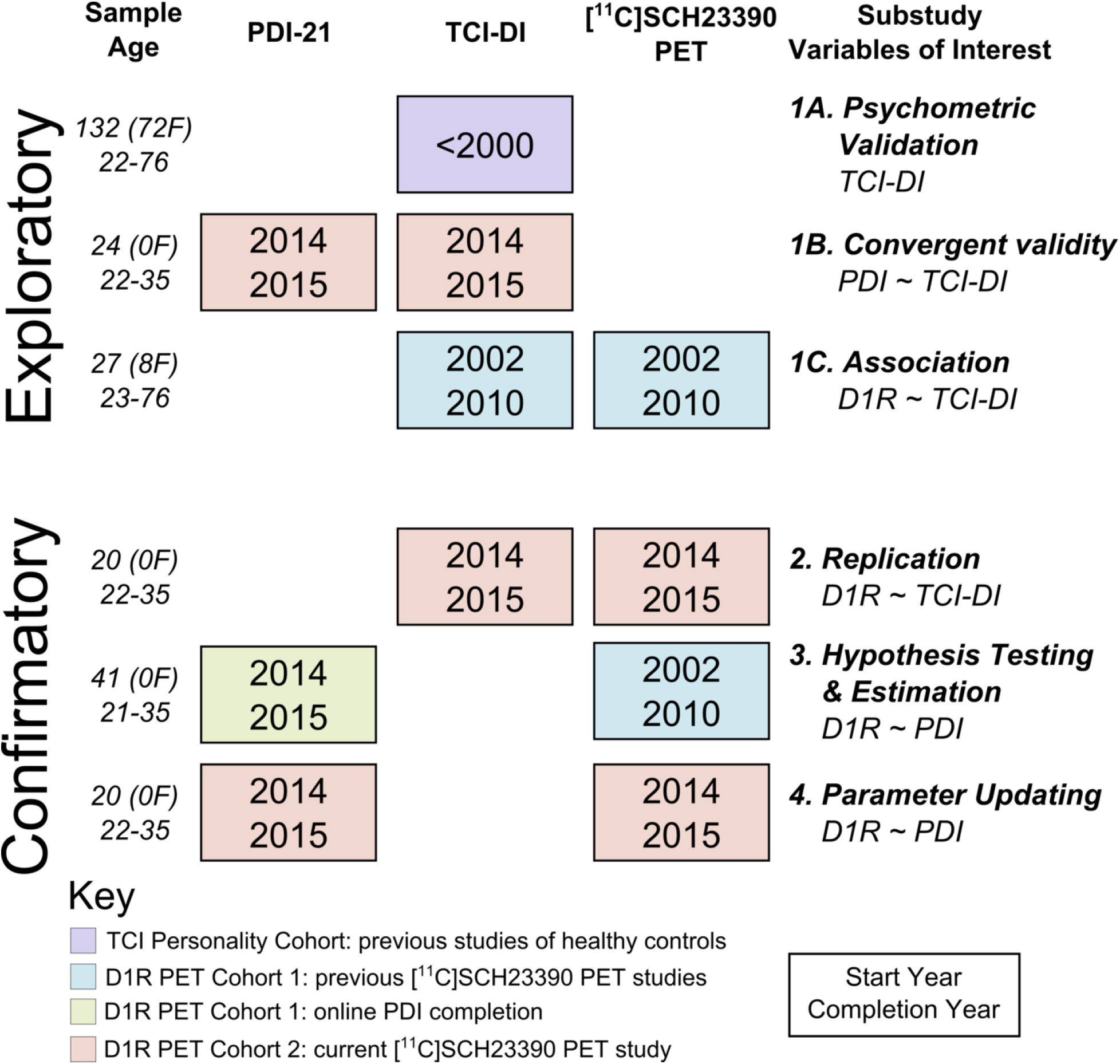
Summary of the included substudies.

### Participants

#### TCI Personality Cohort

132 subjects (72 women) completed the Temperament and Character Inventory (TCI) questionnaire prior to the year 2000. All subjects participated as controls in candidate gene studies conducted at the Department of Psychiatry at Karolinska Institutet. Participants were aged 47.9 ± 16.1 (mean ± SD).

#### D1R PET Cohort 1

All subjects in this cohort participated as control subjects in studies conducted at the Karolinska Institutet PET centre between 2002 and 2010. In all studies, participants underwent physical and mental health examinations before participation, and were judged to be healthy and to show no history of significant psychiatric or somatic illness.

27 participants in this cohort completed the TCI questionnaire at the time of PET. Of these, 20 were younger than 35 years (which we define as ‘young’), of whom 13 were males. Of the participants who did not fall within the ‘young’ age range, all were aged between 69 and 77 years old.

Healthy male participants from D1R PET Cohort 1, who were between the ages of 20 and 35 at the time of PET, were contacted by post in 2014 and 2015 and asked to complete the PDI scale through an online form. This limited age range was selected due to the known influence of age on both [^11^C]SCH23390 BP_ND_ (Wang et al., 1998; Jucaite, Forssberg, Karlsson, Halldin, & Farde, 2010; Bäckman et al., 2011) and PDI scores(Peters, Joseph, Day, & Garety, 2004; Fonseca-Pedrero, Paino, Santarén-Rosell, Lemos-Giráldez, & Muñiz, 2012), as well as the bimodal distribution of age within the sample. Of the 55 participants contacted, 41 participants completed the questionnaire, all of whom are included in the current analysis. Included participants were aged between 20.7 and 35.3 years (mean: 25.0, SD: 2.9) at the time of PET. The number of years between PET and questionnaire completion was 7.9 ± 1.6 (mean ± SD) (range: 4.8 − 12.7 years). The original quantitative analysis of the PET data (i.e. BP_ND_ values) was not used for this study. Instead, the data were re-analysed using the same processing pipeline as used for D1R PET Cohort 2 which was collected more recently.

#### D1R PET Cohort 2

Twenty nine subjects participated in a study conducted at the Karolinska Institutet PET centre between 2014 and 2015. All participants underwent an examination of medical history, physical examination, Magnetic Resonance Imaging (MRI), blood and urine chemistry test, psychiatric screening based on the M.I.N.I. interview, Becks Anxiety Inventory, the Montgomery-Åsberg Depression Rating Scale, and the Alcohol Use Disorders Identification Test. Following these assessments, participants were deemed to be healthy and not to show any history of significant psychiatric or somatic illness.

Of these participants, 22 underwent two PET measurements to examine the test-retest reliability of [^11^C]SCH23390 (Matheson et al., 2017). PET images from the first six of these participants were excluded due to technical issues with the radioligand synthesis. Seven additional participants underwent only one PET examination. Of the 23 subjects with at least one valid PET measurement, PET measurements from two individuals were excluded from the analysis due to incorrect placement in the gantry resulting in only a small portion of the cerebellum being visible. In addition, one PET measurement was excluded due to technical failure of the PET system, resulting in missing frames at the start of the measurement. One participant erroneously left one page of their PDI questionnaire blank, and was therefore excluded from analysis of the PDI. One participant did not complete either questionnaire and was excluded from this study. This resulted in a final sample size of 20 PET measurements, 28 TCI questionnaires, and 27 PDI questionnaires. Included participants were aged between 21.8 and 35.0 years (mean: 25.9, SD: 3.2).

#### Ethics

All studies were approved by the Regional Ethics Committee in Stockholm and all subjects provided informed consent prior to their participation in the studies. All studies making use of PET imaging were approved by the Radiation Safety Committee of the Karolinska Hospital.

### MRI Procedures

T1- and T2-weighted MRI images were acquired for all participants. Within D1R PET Cohort 1, these were measured using a 1.5T GE Signa system (Milwaukee, WI). Within D1R PET Cohort 2, these were measured using a 1.5T Siemens Magnetom Avanto system (Erlangen, Germany). T2-weighted images were examined for structural pathology as a criterion for subject inclusion. The T1-weighted images were used for all further data processing and delineation of regions of interest (ROIs).

### Radiosynthesis and PET Experimental Procedure

Plaster helmets were made for all PET participants, and used during the PET measurements to minimise head movement (Bergstrom et al., 1981). All PET measurements were conducted using an ECAT Exact HR 47 system (CTI/Siemens, Knoxville, TN) run in 3D mode. Transmission scans were performed prior to each PET measurement using three ^68^Ge rods to correct for signal attenuation.

[^11^C]SCH23390 was prepared from [^11^C]methyl iodide (Halldin et al., 1986) and injected into the antecubital vein as a rapid bolus. Injected radioactivity ranged between 221 and 350 MBq (mean: 304, SD: 25 MBq) in cohort 1, and between 248 and 417 MBq (mean: 336, 336, SD: 46 MBq) in cohort 2. Data were acquired for 51 min in a consecutive series of thirteen time frames of durations 3×60 s, 4×180 s and 6×360 s. Following correction for attenuation, random and scattered events, images were reconstructed using filtered back projection, with a Hann filter of 2 mm cut off frequency. The reconstructed volume was displayed as 47 horizontal sections with a voxel size of 2.02 mm × 2.02 mm ×3.125 mm.

### Image Preprocessing

PET images were corrected for residual head motion using post-reconstruction frame-by-frame realignment in which all frames were individually aligned to the first minute of acquisition using SPM5 (Wellcome Department of Cognitive Neurology, University College London). MR images were first reoriented into the AC-PC plane, and then coregistered to the PET images using SPM5.

### Region of Interest Delineation and Quantification

For the delineation of the dorsolateral prefrontal cortex (DLPFC), we used the HCP MMP 1.0 atlas (Glasseret al., 2016) projected onto fsaverage space (Mills, 2016). From this atlas, we combined the following regions: 9-46d, 46, a9-46v, and p9-46v to obtain a region principally consisting of Brodmann’s Area 46. We aimed to select a large enough region for reliable quantification, but specific enough to represent the DLPFC, which we considered the most relevant region for the research question (Wagstyl et al., 2016; Cannon et al., 2002; Selemon & Goldman-Rakic, 1999; Callicott et al., 2000; Abi-Dargham et al., 2002; Abi-Dargham et al., 2012). These regions were merged, converted to individual MR space, and coregistered to the PET images.

For the delineation of the striatum, we used the FSL maximum probability Oxford-Imanova Striatal Connectivity Atlas with three subdivisions thresholded at 25% (Tziortzi et al., 2014). We combined and binarised all three regions to create a striatum ROI in MNI space, and multiplied it by the FreeSurfer grey matter segmentation mask to select only grey matter voxels. The striatum was included as a region with higher reliability (Hirvonen,Nagren, Kajander, & Hietala, 2001; Matheson et al., 2017; Kaller et al., 2017), but lower face validity for the research objective than the DLPFC.

The cerebellar grey matter was used as a reference region as it contains negligible levels of D1R (Chan et al., 1998). The ROI was defined using the maximum probability FSL MNIfnirt cerebellar atlas segmentation with a 25% threshold (Diedrichsen, Balsters, Flavell, Cussans, & Ramnani, 2009), from which a mask was defined and trimmed such that it came no closer than 8mm from the cortex, 8mm from the vermis and 4mm from the edge of the brain. Next, voxels were excluded which belonged to the two most inferior planes of the PET image, and which were identified as belonging to the grey matter using the FreeSurfer segmentation mask. For more details, see (Matheson et al., 2017).

Time activity curves were extracted from all ROIs and regional binding potential (BP_ND_) values were estimated using the simplified reference tissue model (SRTM) (Lammertsma & Hume, 1996) with cerebellar grey matter as the reference region using the R package *kinfitr (v 0.2.0) (Matheson, 2017)*.

### Psychometric Assessment

A Swedish translation of the 21-item Peters Delusion Inventory (PDI) (Peters, Joseph, Day, & Garety, 2004) and the Swedish language version of the 238-item Temperament and Character Inventory (TCI)(Brandstrom et al., 1998; Cloninger, 1993) were used in this study. The Self-Transcendence (ST) scale of the TCI was originally intended to measure creativity and spirituality (Cloninger, 1993), but has later been shown to be associated with proneness to psychosis, as well as clinical psychotic phenotypes (Smith, Cloninger, Harms, & Csernansky, 2008; Daneluzzo, Stratta, & Rossi, 2005; Cortés et al., 2009; Guillem, Bicu, Semkovska, & Debruille, 2002; de Padilla etal., 2006; Glatt, Stone, Faraone, Seidman, & Tsuang, 2006). In order to specifically examine delusional ideation, we created the new scale used in this study by selecting TCI items from the ST scale which exhibit face validity for this construct. Psychometric validation was performed using the R package *psych* (Revelle, 2017). For both inventories, missing items were corrected for by calculating the average score on each item of the scale, and multiplying this by the number of questions.

### Statistical Analysis

All statistical analyses were performed using R version 4.2 (‘Short Summer’)(R Core Team, 2017).

#### Exploratory Analyses

##### Psychometric analysis (Substudies 1A-B)

For Substudy 1A, in which TCI-ST items were selected to create a new measure of delusional ideation, we used Cronbach’s α to assess split-half reliability. We considered a reliability of about 0.7 to be acceptable, according to the heuristic recommendation by (Nunnally & Bernstein, 1978), as the present investigation is considered to be basic research. For Substudy 1B, we assessed the convergent validity of the TCI-DI scales by comparing TCI-DI scores with the more well-established PDI questionnaire within the same individuals, using Pearson’s correlation coefficient.

##### Exploratory correlational analysis (Substudy 1C)

To compare TCI-DI scores with BP_ND_, we made use of multiple linear regression taking the effects of the nuisance variables of age and gender into account. Due to the bimodal age distribution and the highly unequal gender ratio in this data set, there was little reason to prefer exclusion of older individuals and/or women, or inclusion of these variables in the regression model. Furthermore, this analysis was undertaken prior to the collection of the data from Substudy 1B, and therefore made use of both the longer and shorter variants of the TCI-DI scale. Due to the multiplicity of potential analysis decisions which could potentially have been undertaken for this analysis, we made use of a multiverse analysis (Steegen, Tuerlinckx, Gelman, & Vanpaemel, 2016). This means that we present the results following each potential analysis decision, i.e. each path within the “ garden of forking paths” (Gelman & Loken, 2014), with regard to subject inclusion and/or correction for nuisance variables, and draw conclusions based on the general distribution of outcomes. This helps to guard against the influence of hindsight bias in cherry-picking the most optimistic outcomes, thereby increasing the risk of type I errors.

#### Confirmatory Analyses

##### Bayesian Statistical Analysis

For the confirmatory analyses, we made use of Bayesian inference. This means that statistical parameters, such as regression coefficients, are represented by probability distributions which represent relative degree of belief in different values for that parameter. Observed data are then used to update our *prior* information or beliefs about the probability of different values of the parameter, to arrive at a *posterior* distribution (Lee & Wagenmakers, 2014). This is referred to as Bayesian parameter estimation.

We also performed Bayesian hypothesis testing, in which we compare the predictive adequacy of competing models representing different hypotheses about the data with one another, giving rise to the Bayes Factor (BF). The BF is the extent to which the data update the prior model odds of the two models to the posterior model odds. A BF_12_ of 5 means that the observed data are 5 times more likely to have occurred under model 1 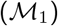 compared to model 2 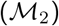. As such, the BF represents the relative probability of the observed data under one hypothesis compared to another (Lee & Wagenmakers, 2014; Ly, Verhagen, & Wagenmakers, 2016). Hence, it signals how much support, or evidence, there is for one hypothesis over the other.

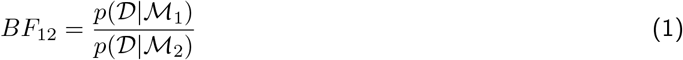

In this way, different models can also be compared indirectly as follows:

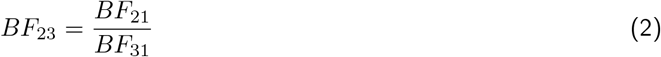

BFs were calculated using the Savage-Dickey density ratio method with a point null hypothesis (Dickey, 1971).

##### Replication Analysis (Substudy 2)

For performing a replication analysis of the results of Substudy 1C, we made use of the correlation replication BF (Wagenmakers, Verhagen, & Ly, 2016), with the posterior calculated analytically (i.e. without the need for sampling). The correlation replication BF assumes that the original finding was obtained using a flat (uniform) prior. The posterior of this analysis is used as the prior for the replication attempt, which is compared to a null hypothesis. In this way, the replication BF can be thought of as a comparison of the skeptic’s null hypothesis with the proponent’s posterior of the original findings, and evaluates the relative probability of these two hypotheses given a new set of data. Put differently, the replication BF represents the relative change in likelihood of the estimated effect being equal to 0 before, and after, having observed the data from the replication study (Wagenmakers, Verhagen, & Ly, 2016).

##### Multiple Regression Analysis (Substudies 3-4)

For substudies 3 and 4, we made use of Bayesian multiple linear regression to examine the association between PDI scores and [^11^C]SCH23390 BP_ND_. The regression model was specified using unstandardised units. This restricts the potential for limited variation in delusional ideation scores to influence the parameter of interest.

We defined informative priors derived from the literature to account for the effects of age on BP_ND_ and PDI scores, a uniform prior for the intercept, and we defined zero-centred regularising priors for the parameter of interest. The definition of the priors is described in Supplementary Materials S4. The model was specified as follows:

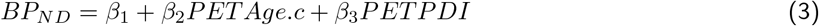

PET.Age.c represents the mean-centred age at the time of PET. PETPDI represents the individuals’ theoretical PDI score when they were at the age around which PETAge is centred (i.e. the mean age of all participants at the time of PET). Individuals’ PETPDI scores were estimated as follows:

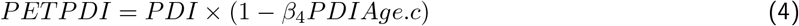

where PDI represents raw PDI scores, and PDIAge.c represents participants’ age at the time of PDI completion centred to the mean PET age. For Substudy 3, PDIAge.c is greater than PETAge.c to account for the years elapsed between PET and PDI completion. For Substudy 4, they are the same value, as all participants completed the questionnaire within two weeks of PET. We can therefore derive a final model by substituting equation 3 into equation 4:

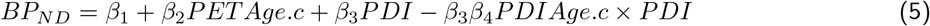

Model fitting in studies 5 and 6 was performed using Gibbs Markov Chain Monte Carlo (MCMC) sampling using JAGS (Plummer, 2003) and the R package *R2jags (Su & Yajima, 2015)*.

##### Bayesian Hypothesis Testing

In Substudy 3, we made use of Bayesian hypothesis testing. Based on the mixed results of clinical PET studies comparing D1R availability in schizophrenia patients and controls (Okubo et al., 1997; Abi-Dargham et al., 2002; Karlsson, Farde, Halldin, & Sedvall, 2002; Hirvonen et al., 2006; Kosaka et al., 2010; Abi-Dargham et al., 2012; Poels, Girgis, Thompson, Slifstein, & Abi-Dargham, 2013), we defined and compared three hypotheses: an increase, a decrease, and no change in D1R BP_ND_ with increasing proneness to psychosis.

In order to generate statistical definitions of these hypotheses, we assumed that the differences between the mean [^11^C]SCH23390 BP_ND_ and mean PDI scores between patient and control groups would correspond with one another between studies. For PDI scores, (Peters, Joseph, Day, & Garety, 2004) observed a difference of five points between delusional patients and controls. For DLPFC [^11^C]SCH23390 BP_ND_, the effect magnitude was defined separately for each of the three hypotheses. More details are provided in Supplementary Material S4.

##### Bayesian Parameter Estimation

In Substudies 3 and 4, we performed Bayesian parameter estimation to investigate the association between D1R BP_ND_ and PDI scores. This method makes weaker assumptions about the magnitude of the relationship than the Bayesian hypothesis testing method above, but yields conclusions relating to the magnitude and direction of the effect rather than the relative likelihood of the different hypotheses. For Substudy 3, we used the widest of the hypothesis testing priors above as a bidirectional regularising prior to estimate the strength of association. For Substudy 4, we used mean-scaled normal approximations of the posterior estimates from Substudy 3 as priors for all parameters except for the intercept in order to update posterior estimates in an independent sample.

### Data and Code Availability

Analysis code, and data which is not considered to be sensitive, will be made available at https://github.com/mathesong/D1R_PsychProneness. The data which are not publicly available due to regulatory restrictions are available upon request from the authors.

## Results

### Exploratory Analysis

#### Substudy 1: Exploratory analysis of the Association between Delusional Ideation and D1 receptor BPND measured using [^11^C]SCH23390

In order to assess the relationship between delusional ideation and D1R availability using previously collected data, we selected items from the TCI-ST scale which were considered to be representative of delusional ideation. Using these items, we created a short and a long TCI Delusional Ideation (TCI-DI) scale. These scales were shown to exhibit good to acceptable reliability (Cronbach’s *α* = 0.82 and 0.76 respectively) (Supplementary Materials S1). The scales were both positively associated with the PDI in an independent sample, thereby demonstrating convergent validity. The short version of the TCI-DI scale displayed a numerically higher correlation compared to the long version (r=0.64 vs r=0.46) (Supplementary Materials S2). The short TCI-DI scale was therefore considered to be the primary TCI-DI scale.

Next, we assessed the relationship of these scales with [^11^C]SCH23390 BP_ND_ in the DLPFC and striatum using a multiverse analysis. We observed a negative relationship between TCI-DI scores and [^11^C]SCH23390 BP_ND_ of generally large effect size (Figure 2). This association did not appear to be greatly influenced by the different analytical alternatives, although effect sizes tended to be larger when examining only young male participants rather than including the whole sample with gender and age as covariates in the model. More details are provided in Supplementary Materials S3.

**Figure 2:**
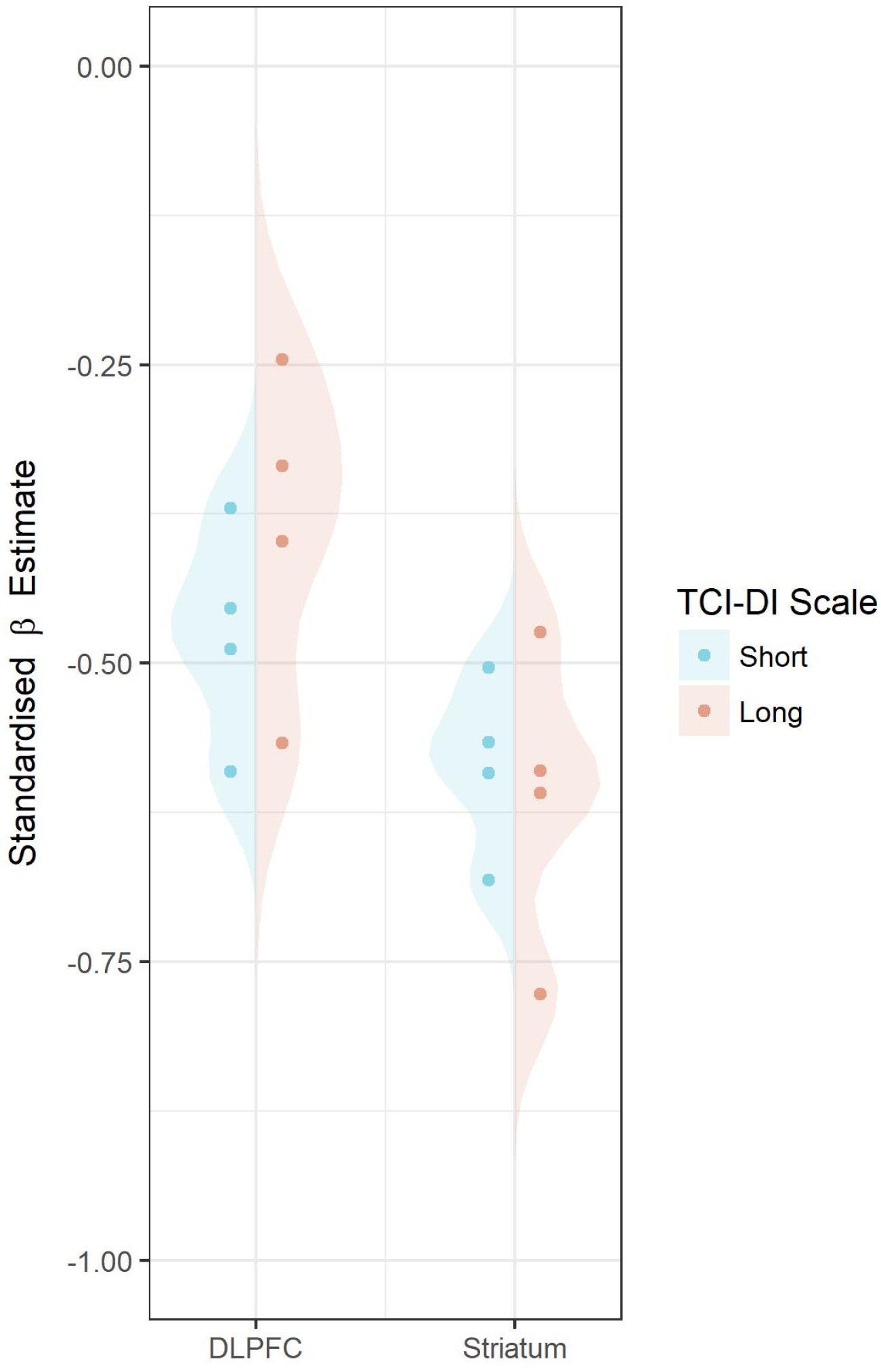
Distribution of the effect sizes from the multiverse analysis for the association between TCI-DI scores and D1R availability following different analytical paths.

In summary, contrary to our hypothesis of a positive association, we found preliminary evidence of a negative association between delusional ideation and D1R availability in both the DLPFC and striatum. We therefore considered the association between delusional ideation and D1R availability a promising hypothesis to be tested in a confirmatory study.

### Confirmatory Analyses

#### Substudy 2: Confirmatory replication of results from Substudy 1C

We examined the relationship between [^11^C]SCH23390 BP_ND_ and TCI-DI scores in young males to assess whether the results of Substudy 1C could be replicated in an independent sample. The Cronbach’s α of the TCI-DI scale in this sample was 0.61. In contrast to the results of Substudy 3, there was no apparent association between TCI-DI scores and BP_ND_ (Figure 3). Replication BFs showed moderate to strong support in favour of the null hypothesis relative to the posterior distribution of the correlation coefficient from the original study (DLPFC BF_0R_=5.5, Striatum BF_0R_=10.5).

**Figure 3:**
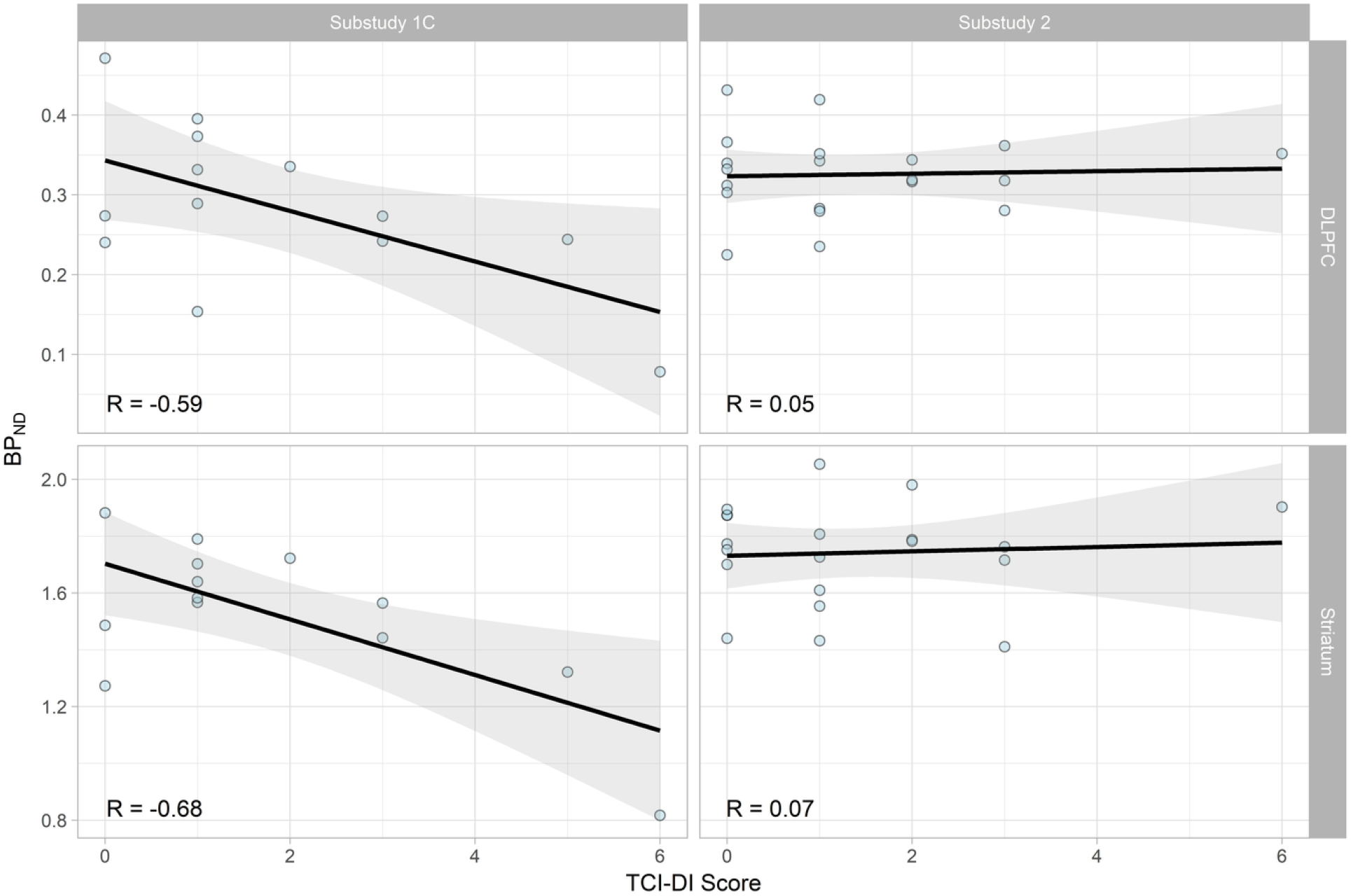
Scatterplots demonstrating the association and their effect sizes between TCI-DI scores and [^11^C]SCH23390 BP_ND_ in the DLPFC and striatum from the cohorts examined in Subsubstudies 3 and 4.

In summary, this means that the data were five to ten times more likely under the null hypothesis of no association compared to that of the negative association seen in Substudy 1C.

#### Substudy 3: Confirmatory analysis of the association between PDI scores and D1 receptor BP_ND_ measured using [^11^C]SCH23390

This sample consisted of participants who had previously undergone PET measurements with [^11^C]SCH23390, who later completed the PDI through an online questionnaire. In this sample, the Cronbach’s α for the PDI was 0.78. The number of years elapsed between participants’ PET measurements and their completion of the scale ranged between 4.8 and 12.7 years (mean 7.9, sd 1.6). Despite the limited variation in age in the data set, we still observed negative correlations with age for both DLPFC BP_ND_ (r = −0.18) as well as PDI scores (r =-0.23) as expected from previous literature (Wang et al., 1998; Jucaite, Forssberg, Karlsson, Halldin, & Farde, 2010; Bäckman et al., 2011; Fonseca-Pedrero, Paino, Santarén-Rosell, Lemos-Giráldez, & Muñiz, 2012; Peters, Joseph,Day, & Garety, 2004). Informative priors derived from the literature were defined to account for the effects of age on both D1R BP_ND_ and PDI scores in the model.

##### Bayesian Hypothesis Testing

Bayes Factors (BFs) comparing the relative likelihood of the data under each of the three hypotheses describing the association between D1R BP_ND_ and delusional ideation are shown in Table 2.

We found medium to strong evidence in favour of the null hypothesis compared to the other two hypotheses. This means that the data is over 5 times more likely to have occurred under the null hypothesis model than it is under the increase model, and over 10 times more likely under the decrease model.

We also performed a similar analysis for the striatum, assessing the hypothesis of a negative association between striatal BP_ND_ and PDI scores based on the results of Substudies 1B and 1C. In this analysis, we found strong support for the null hypothesis (BF_01_ = 17.9), supporting the results of Substudy 2. For this reason, and since the DLPFC was our primary region of interest, the striatum was not further examined in Substudy 4.

##### Bayesian Parameter Estimation

The posterior density for the association between PDI scores and BP_ND_ values (Beta3), was found to be concentrated close to zero for both the DLPFC and the striatum (see Figure 4 for the DLPFC, and Supplementary Materials S5 for the striatum). This corresponds to changes in [^11^C]SCH23390 BP_ND_ of 1.8% (95% CredInt: −5.8 − 9.6%) for the DLPFC and 3.3% (95% CredInt: −3.3 − 6.5%) for the striatum, for an increase of 5 points on the PDI scale. Posterior summaries are provided in Supplementary Materials S6.

**Table 1:** Bayes Factors comparing each hypothesis (rows) against each other hypothesis (columns)

| Model | Decrease | Increase | Null |
| --- | --- | --- | --- |
| Decrease | 1.00 | 0.55 | 0.10 |
| Increase | 1.83 | 1.00 | 0.18 |
| Null | 10.45 | 5.69 | 1.00 |

**Figure 4:**
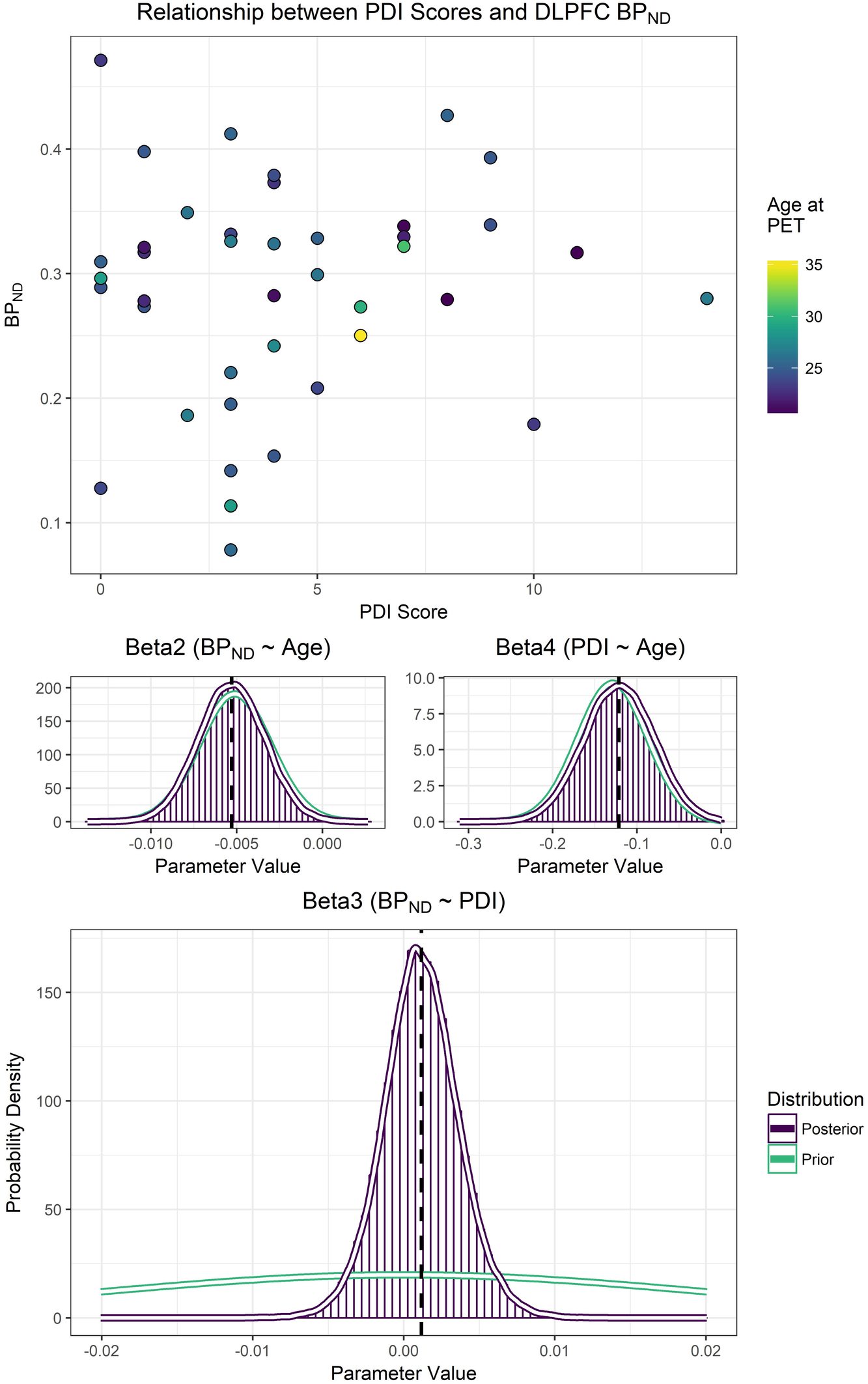
Scatterplot (top) displaying the association between PDI scores and BP_ND_ values in DLPFC for Substudy 3. Below are posterior density plots for the unstandardised beta estimates of the regression. In the middle row are the parameters representing the effects of age on BP_ND_ (left), and PDI scores (right). In the bottom row is the parameter representing the change in BP_ND_ values for each increase in PDI score.

#### Substudy 4: Bayesian updating of the results of Substudy 3

We aimed to update parameter estimates from Substudy 3 in an independent sample of individuals who had completed PDI scales within two weeks of the PET measurements. The Cronbach’s α of the PDI scale in this sample was 0.68. Estimates for the association between DLPFC [^11^C]SCH23390 BP_ND_ and PDI scores were calculated using the same regression model as in Substudy 3, using posterior estimates from Substudy 3 scaled to the mean BP_ND_ in this sample as priors for all parameters except for the intercept (see Supplementary Materials S4).

The posterior distribution for the regression coefficient representing the association between PDI scores and DLPFC BP_ND_ (Beta3) was updated such that it became both more precise, but also closer to zero (Figure 5). According to this final estimate, a change of 5 points on the PDI, the margin of difference previously observed between populations of healthy controls and delusional patients(Peters, Joseph, Day, & Garety, 2004), is associated with a change in BP_ND_ of 1.5% of the mean (95% CredInt: −6.1 − 7.7%).

**Figure 5:**
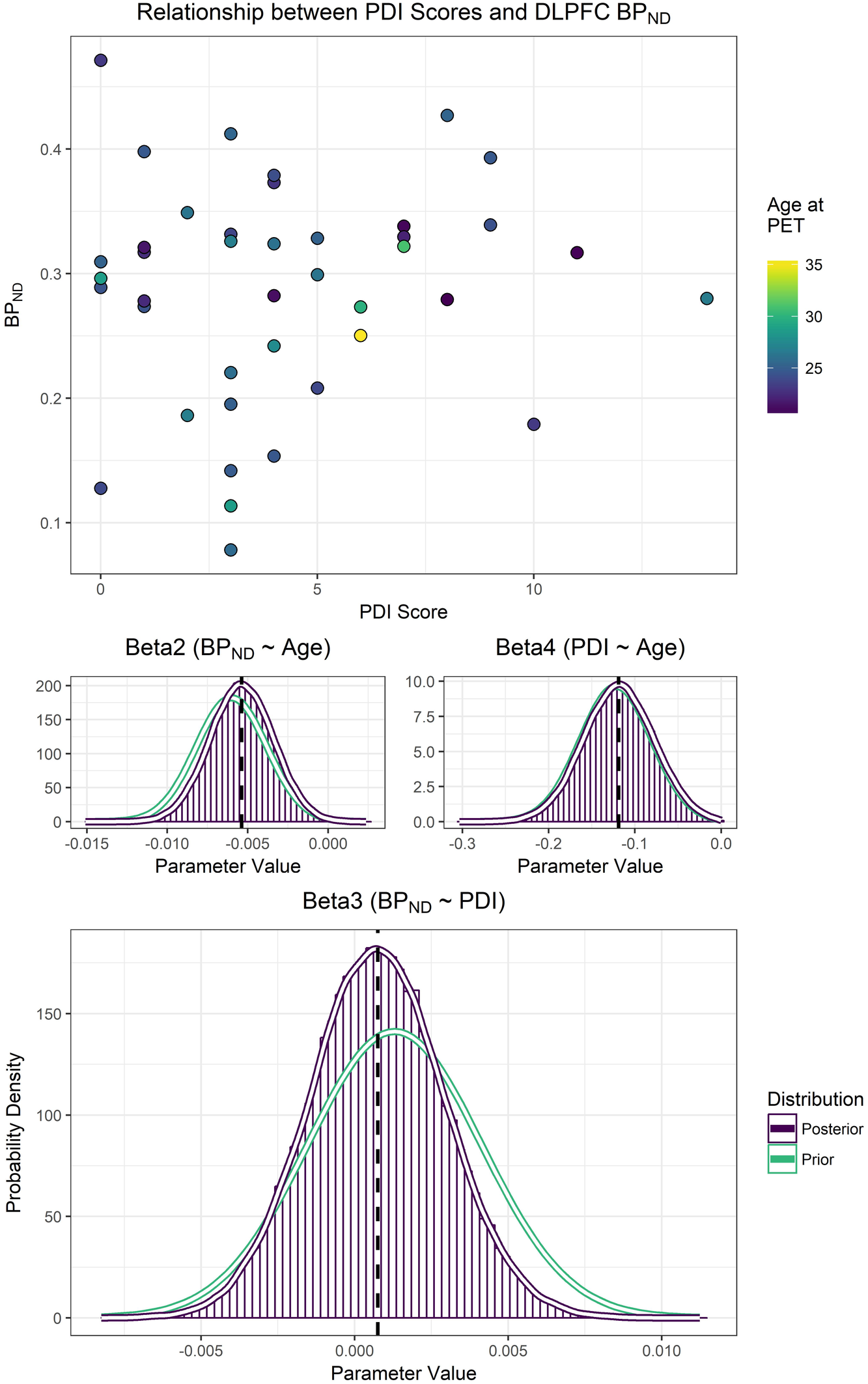
Scatterplot (top) displaying the association between PDI scores and BP_ND_ values in DLPFC for Substudy 4. Below are posterior density plots for the unstandardised beta estimates of the regression using the posterior estimates from Substudy 3 as priors. In the middle row are the parameters representing the effects of age on BP_ND_ (left), and PDI scores (right). In the bottom row is the parameter representing the change in BP_ND_ values for each increase in PDI score.

## Discussion

In this study, we aimed to investigate the relationship between D1R availability and delusional ideation using a combination of exploratory and confirmatory analyses, in order to generate and test hypotheses respectively. In the exploratory part of the investigation, we created a new scale for measurement of delusional ideation from items of the TCI questionnaire and validated its psychometric properties (Substudies 1A and 1B). This scale exhibited a negative association with dopamine D1R availability in both the DLPFC and striatum (Substudy 1C). In the confirmatory part of this investigation, we first found moderate to strong evidence that the results of Study 1C could not be successfully replicated, suggesting that they were more likely to be a false positive (Substudy 2). We went on to demonstrate evidence in favour of no association between PDI scores and D1R availability (Substudy 3). Further estimation also revealed that the expected changes in D1R availability for increasing scores on the PDI scale were too small to be of relevance (Substudies 3-4). In summary, our results show, with a high degree of precision, that there is little to no linear relationship between DLPFC D1R availability and delusional ideation in healthy controls.

Studies examining DLPFC [^11^C]SCH23390 BP_ND_ in schizophrenia patients and controls have found a 27% reduction(Kosaka et al., 2010), a 35% increase(Poels, Girgis, Thompson, Slifstein, & Abi-Dargham, 2013), a 26% reduction (Hirvonen et al., 2006) and no difference (5%) (Karlsson, Farde, Halldin, & Sedvall, 2002) compared to controls, as well as a 21% increase in monozygotic twins of schizophrenia patients compared to controls(Hirvonen et al., 2006). Based on the final parameter estimates from the present investigation, a change in 5 points on the PDI scale, previously found to be the difference between delusional patients and controls (Peters, Joseph, Day, & Garety, 2004), was only associated with a 1.5% change in BP_ND_ (up to 7.7% within the 95% Credible Interval), which is not consistent with the results of any of the previous studies reporting significant differences between groups. To contextualise this result further, the resulting model from Substudy 4 estimates that a change across the entire extent of the PDI scale, 21 points, would still only amount to a 6.5% change in BP_ND_ (−25.8 − 32.3%: 95% Credible Interval). Although the credible interval now theoretically overlaps with some previous estimates of the differences between patients and controls, such a large difference in PDI scores is unrealistic. We can therefore conclude that our results are not consistent with the magnitude of change which would have been expected based on studies examining patients and controls.

**Table 2:** Posterior summaries of unstandardised coefficients from updated parameters

| Parameter | mean | sd | 2.5% | 97.5% | Rhat |
| --- | --- | --- | --- | --- | --- |
| Beta1 | 0.323 | 0.015 | 0.293 | 0.352 | 1 |
| Beta2 | -0.005 | 0.002 | -0.009 | -0.002 | 1 |
| Beta3 | 0.001 | 0.002 | -0.004 | 0.005 | 1 |
| Beta4 | -0.119 | 0.041 | -0.200 | -0.039 | 1 |

The interpretation of cortical [^11^C]SCH23390 and [^11^C]NNC112 binding is limited by the fact that both radioligands are sensitive to binding to serotonin 2A receptors(5-HT_2A_R), estimated to account for approximately 25% of cortical binding for both radioligands (Ekelund et al., 2007). Studies suggest that 5-HT_2A_R availability may be decreased in schizophrenia patients compared to controls (Rasmussen et al., 2016; Rasmussen et al., 2010; Ngan, Yatham, Ruth, & Liddle, 2000) (although some older studies did not observe significant differences: (Lewis et al., 1999; Okubo et al., 2000; Trichard et al., 1998)), and several antipsychotic drugs have an affinity for the 5-HT_2A_R (Schotte et al., 1996; Correll et al., 2015). It is therefore possible that both previous and present results may have been affected by differences in 5-HT_2A_R availability. If both D1R and 5-HT_2A_R availability differ with increasing proneness to develop schizophrenia, and if these changes are of opposite sign, then this could result in the present findings of no effects despite a true underlying change. This issue could be addressed by the assessment of the association between delusional ideation and 5-HT_2A_R availability. Two preliminary studies with small sample sizes have examined this, but the results have not been consistent (Rasmussen et al., 2016; Hurlemann et al., 2008).

A limitation of our study was the time elapsed between PET and completion of PDI questionnaires in Substudy 3. However, based on the consistency of the results of Substudy 3 with those of Substudy 4 for which there was little to no delay, as well as the close correspondence of the literature prior and posterior densities for Beta4, the influence of this limitation appears to have been successfully accounted for by the model. A second limitation of this study is the restricted inter-individual variation in delusional ideation scores in our sample (Supplementary Materials S7). We observed that the PDI scores were right-skewed, which has been previously reported(Peters,Joseph, Day, & Garety, 2004; Peters, Joseph, & Garety, 1999). In Substudies 3 and 4, however, we made use of Bayesian linear regression without variance scaling (i.e. using unstandardised units). Standardised coefficients, such as the correlation coefficient, express a relationship between variables relative to the degree of variance in the sample, and will be lower if there is less variance. In contrast, our use of unstandardised units allowed us to demonstrate that the probability of there being a meaningfully large change in BP_ND_ with increasing levels of delusional ideation is essentially nonexistent. In other words, limited variation of scores within the sample is unlikely to have given rise to the observed result of a highly precise posterior distribution centred around zero.

Our analysis suggested strong, convergent evidence for a lack of association between D1R availability and delusional ideation. Several potential interpretations of these results can be considered in relation to previous studies of the D1R in schizophrenia. A first possible interpretation is that there may simply be no association between D1R availability and schizophrenia at all, consistent with the results of (Karlsson, Farde, Halldin, & Sedvall, 2002). A second alternative is that changes in D1R availability may only occur at the onset of the disorder. A third could be that D1R availability is unrelated to delusional ideation, but rather associated with some other component of proneness to develop psychosis. Psychosis proneness is unlikely to be a unidimensional construct, and D1R changes may instead be associated with specific genetic risk factors, or other behavioural phenotypes such as negative symptoms or cognitive impairment which are thought to be largely independent of positive symptoms in clinical schizophrenia. A final possibility of a more speculative nature, but which cannot be excluded due to the limited variation in our data, is that of there being a nonlinear association between delusional ideation and D1R availability. This would imply little to no association between these variables at low levels of psychosis proneness, but an increasing degree of association with increasing levels of proneness. Further research into the D1R in schizophrenia and in proneness for schizophrenia is warranted to ascertain which of these alternatives is most likely.

## Acknowledgements

We gratefully thank the members of the PET group at Karolinska Institutet, and of the lab group of Predrag Petrovic for their insightful comments and feedback on the results. We thank Erik Jönsson for collection of the TCI data for Substudy 1A. We thank Per Stenkrona for collection of much of the PET data. This study was funded by The Swedish Research Council (523-2014-3467 and K2012-61X-09114-26-4), Hjärnfonden, Stockholm County Council (ALF, 20150482) and Söderström Königska fonden.

## Author Contributions

**Table 3:**
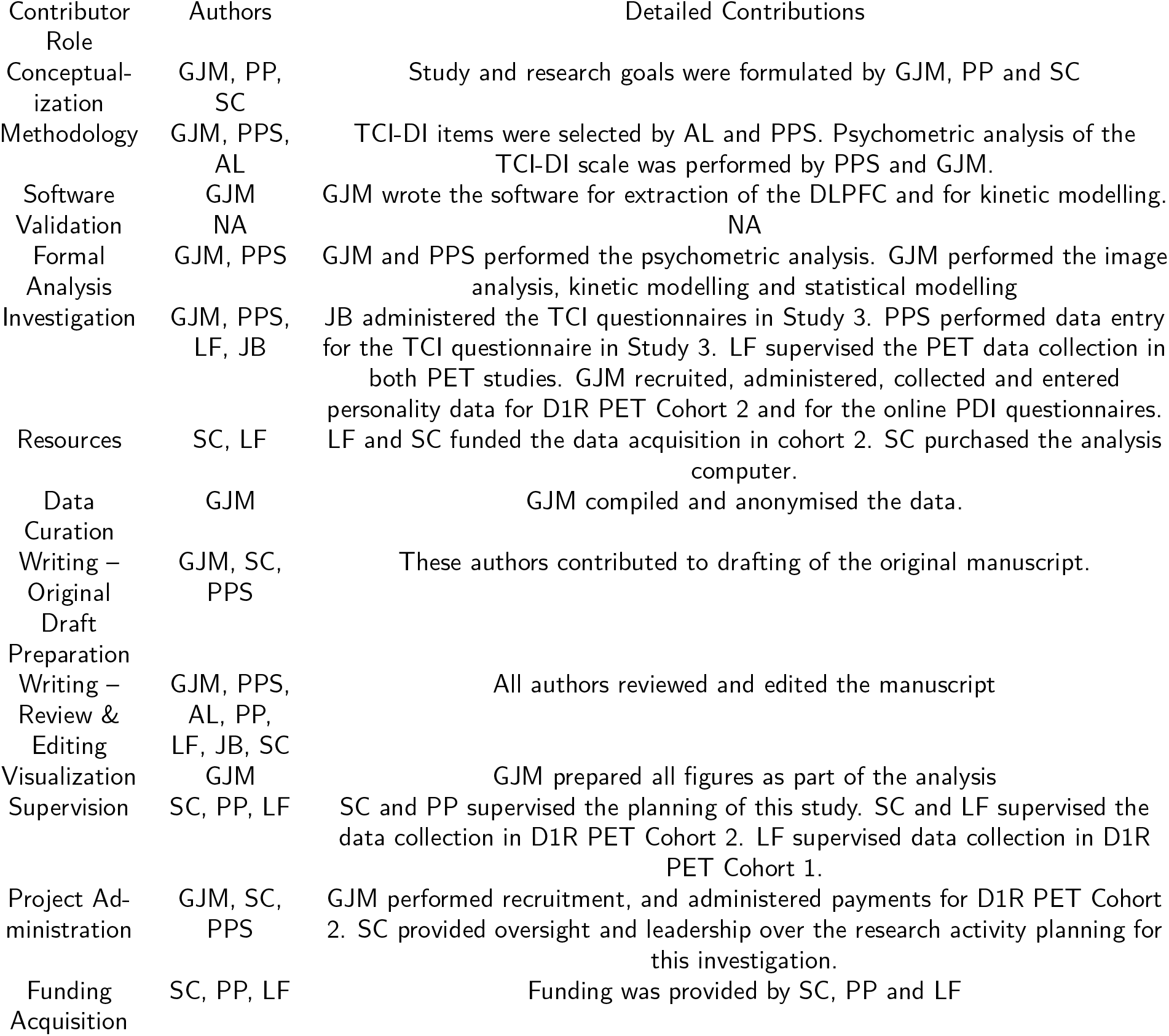
Author Contributions

